# Multiscale volume electron microscopy of the human liver maps vascular-cellular architecture, organelle dynamics and inter-organelle communication

**DOI:** 10.64898/2026.04.22.719970

**Authors:** Cheng Xing, Ronald Xie, Ben Mulcahy, Ali Darbandi, Cornelia Thoeni, Ishaan Singh Chandok, Yuna Lee, Jun Ma, Fez Ali, Ian D. McGilvray, Sonya MacParland, Mei Zhen, Gary D. Bader

**Affiliations:** Department of Molecular Genetics, University of Toronto, Toronto, M5S 1A1, ON, Canada; The Donnelly Centre, University of Toronto, Toronto, M5S 3E1, ON, Canada; Vector Institute, Toronto, M5G 0C6, ON, Canada; NVIDIA, Toronto, M5V 1K4, ON, Canada; Lunenfeld-Tanenbaum Research Institute, Sinai Health System, Toronto, M5G 1X5, ON, Canada; Nanoscale Biomedical Imaging Facility, Hospital for Sick Children, Toronto, M5G 1H3, ON, Canada; Department of Laboratory Medicine and Pathobiology, University of Toronto, Toronto, M5G 3E1, ON, Canada; Department of Physics, Harvard University, Cambridge, 02138, MA, USA; Centre for Brain Science, Harvard University, Cambridge, 02138, MA, USA; Princess Margaret Cancer Centre, University Health Network, Toronto, M5G 2M9, ON, Canada; Multi-Organ Transplant Program, Toronto General Hospital Research Institute, Toronto, M5G 2C4, ON, Canada; Ajmera Transplant Centre, University Health Network, Toronto, M5G 2N2, ON, Canada; Department of Physiology, University of Toronto, Toronto, M5S 3K3, ON, Canada; Department of Computer Science, University of Toronto, Toronto, M5S 2E4, ON, Canada; CIFAR Multiscale Human Program, CIFAR, Toronto, M5G 1M1, ON, Canada

**Keywords:** Human liver, Volume electron microscopy, Multiscale mapping, Mitochondria–endoplasmic reticulum contacts, Biliary and sinusoidal architecture, Deep learning segmentation

## Abstract

The human liver depends on multiscale structural organization from vasculature to cells to organelles to perform its diverse metabolic functions. A unified three-dimensional view linking these hierarchical scales in intact human tissue would be useful for better understanding these levels and how they relate to each other. We present a high-resolution volume electron microscopy reconstruction of human periportal liver tissue (152 × 140 × 33 µm^3^ at 8 nm pixel size) acquired by serial block-face scanning electron microscopy. Using a multiscale deep learning approach, we performed automated segmentation across the entire volume, enabling comprehensive annotation of vasculature, cells, and organelles. Quantitative analysis of bile duct architecture revealed coordinated scaling between lumen geometry and cholangiocyte number and size. Sinusoidal capillary branches exhibited distinct structural profiles with differential endothelial coverage. Analysis of 35,790 complete mitochondria identified substantial morphological heterogeneity, with elongated mitochondria displaying preferential endoplasmic reticulum (ER) contacts at narrowing sites, a pattern consistent with ER-mediated fission or fusion. This multiscale reconstruction establishes an ultrastructural reference for the healthy human liver and provides a quantitative framework for investigating hepatic physiology and disease.

## 1 Introduction

The human liver performs essential metabolic functions including detoxification, nutrient processing, and bile production [1]. These diverse processes depend on intricate multiscale organization, where structural hierarchies—from vasculature to cells to organelles—coordinate to maintain tissue homeostasis [2–4]. At the vascular and cellular scale, bile ducts and sinusoidal capillaries establish the architectural framework for hepatic function [2, 5]. Bile ducts, lined by cholangiocytes, collect and transport bile produced by hepatocytes, while sinusoidal capillaries facilitate bidirectional exchange between blood and hepatocytes through fenestrated endothelium. At the subcellular scale, organelles exhibit morphological and spatial organization linked to functional specialization. For example, mitochondria–endoplasmic reticulum (ER) contacts regulate metabolic signaling, calcium homeostasis, and mitochondrial dynamics—processes central to liver physiology and implicated in metabolic disease [4, 6–8].

Despite the importance of these multiscale relationships, a unified three-dimensional and dynamic view of human liver tissue ultrastructure is not available. Previous studies of liver vasculature using light microscopy, optical clearing, or computational modeling have characterized associations between vascular architecture and the surrounding cellular environment [2, 9, 10], but lack the resolution to visualize vascular lumens, individual cells, and their organelles simultaneously. At the subcellular level, studies in mouse liver and cultured cells have established ER involvement in mitochondrial morphology and dynamics [4, 6, 11], yet quantitative analysis of mitochondria–ER contacts in intact human liver tissue has not been performed.

Here we present a high-resolution three-dimensional reconstruction of human periportal liver tissue using serial block-face scanning electron microscopy. The reconstructed volume (152 *×* 140 *×* 33 µm^3^ at 8 nm pixel size) encompasses hierarchical structures spanning tissue, vasculature, cells, and organelles. We developed a deep learning framework for automated multiscale segmentation, enabling comprehensive annotation across the entire volume. Quantitative analysis revealed structural heterogeneity in bile duct lumen–cholangiocyte contacts and sinusoidal capillary branching organization. Analysis of 35,790 mitochondria identified substantial morphological heterogeneity, with elongated mitochondria displaying preferential endoplasmic reticulum contacts at narrowing sites—a pattern consistent with dynamics of ER-mediated fission or fusion. Together, this work establishes a multiscale ultrastructural reference for the human liver periportal region, providing both a morphological foundation and an analytical framework for investigating hepatic physiology and disease.

## 2 Results

### 2.1 3D reconstruction of human liver tissue

The human liver displays intricate spatial organization, particularly within the periportal region (Fig. 1a), which harbors portal tracts and diverse cell types fundamental to hepatic function and disease susceptibility [12]. To investigate it at nanoscale resolution, we performed volume electron microscopy (vEM) on a healthy human liver tissue sample encompassing the periportal region. To identify the appropriate region of interest (ROI), we first performed X-ray micro-computed tomography on a resinembedded tissue block (Fig. 1b). The 3D X-ray image provided orthogonal views of the block (xy, xz, and yz) (Supplementary Fig. 1a-c), enabling assessment of overall tissue integrity, staining uniformity, and localization of the periportal area. To further identify the periportal zone, immunohistochemistry was performed using CYP2A6, a periportal marker enzyme, which revealed the expected periportal-to-pericentral gradient (Fig. 1c). The periportal ROI was then manually defined based on anatomical landmarks, including the portal vein, bile duct, and hepatic artery. Following ROI identification, the tissue block was trimmed to expose the selected periportal region (Fig. 1b). The trimmed block was mounted inside a serial block-face scanning electron microscope, where the block surface was sequentially cut and imaged. Accurate stitching within each section and alignment across serial sections was performed using the Scalable Optical Flow-based Image Montaging and Alignment (SOFIMA) pipeline [13]. Overlapping tiles were assembled into complete two-dimensional images (Supplementary Fig. 1d), and adjacent sections were aligned to maintain tissue geometry and anatomical continuity (Supplementary Fig. 1e). The reconstructed volume has an inplane resolution of 8 nm per pixel and a section thickness of 55 nm, resulting in a continuous vEM volume (Fig. 1d, Supplementary Video 1).

**Fig. 1.**
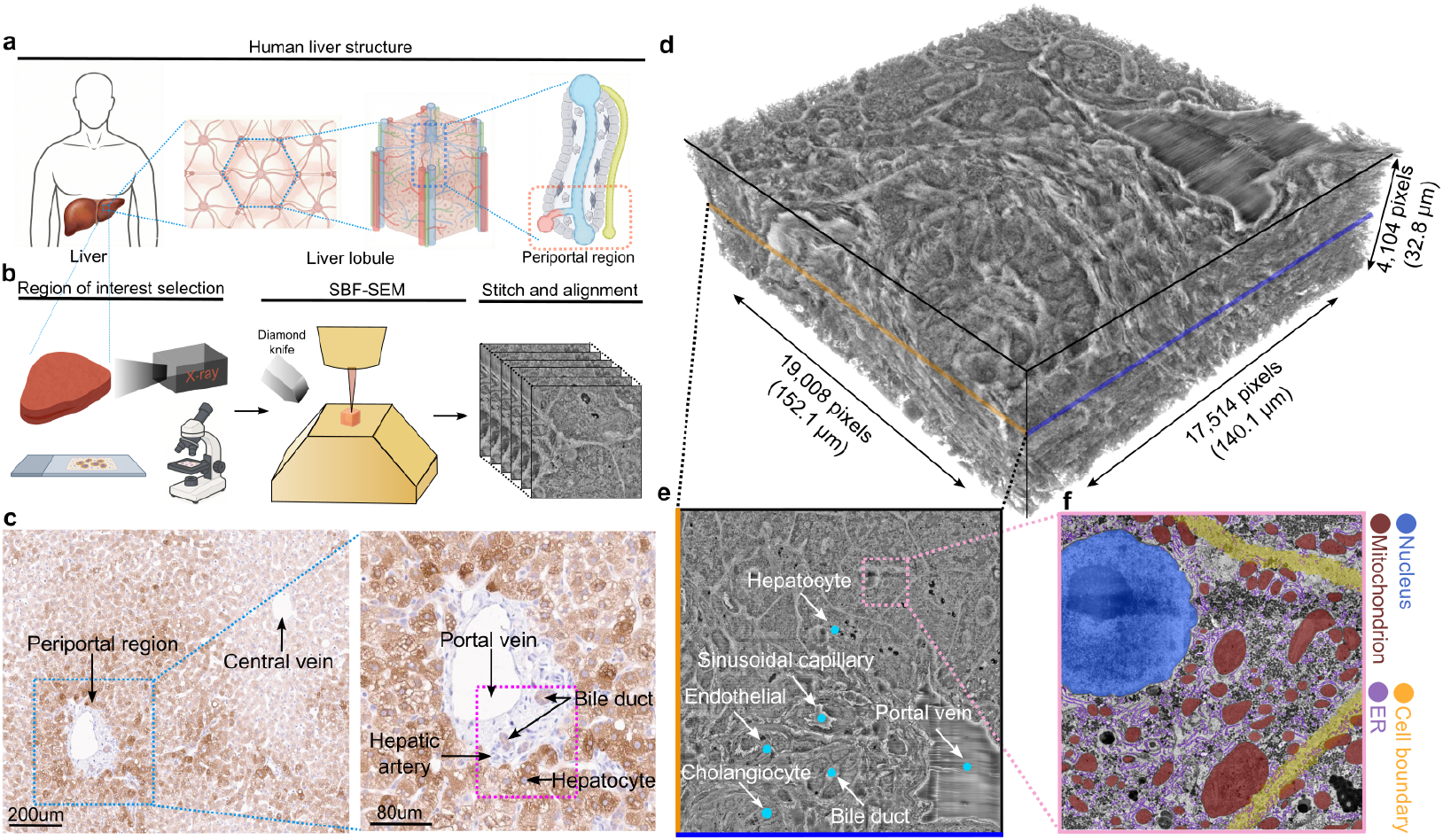
Multiscale volume electron microscopy (vEM) reconstruction of the human liver periportal region. **a**, Schematic illustrating the multiscale organization of liver tissue, from organ to subcellular levels. **b**, Workflow for volumetric reconstruction, including sample preparation, serial sectioning, imaging, and computational assembly. **c**, Left: immunohistochemistry for CYP2A6 demonstrating periportal-pericentral zonation. Right: magnified view of the periportal region showing the portal vein, hepatic artery, bile duct, and surrounding hepatocytes. **d**, Three-dimensional reconstruction of the periportal region showing tissue architecture. **e**, Representative vEM cross-section highlighting cellular and vascular architecture. **f**, Representative vEM cross-section demonstrating subcellular ultrastructure. ER, endoplasmic reticulum.

The periportal region exhibits rich multiscale organization, encompassing vascular networks, multiple cell types, and diverse subcellular structures. To annotate these components, the reconstructed volume was imported into CATMAID [14], a web-based platform for collaborative 3D annotation. An expert pathologist manually annotated vascular and cellular structures by placing coordinate-based markers (Supplementary Fig. 1f). Distinct vascular landmarks including sinusoidal capillaries (Supplementary Fig. 2a), bile ducts, and portal veins were readily identifiable. Major cell types observed include hepatocytes, endothelial cells, and cholangiocytes (Fig. 1e), along with less abundant stellate cells (Supplementary Fig. 2b) and macrophages (Table 1). At the subcellular level, nuclei, mitochondria, endoplasmic reticulum (ER), and cell boundaries can be clearly visualized (Fig. 1f).

**Table 1.** Pathologist-annotated structures and their counts in the multiscale vEM volume.

| Structure | Count |
| --- | --- |
| *Biliary system* |  |
| Bile duct lumen | 1 |
| Bile canaliculi | 1 |
| Cholangiocyte | 143 |
| *Sinusoidal vasculature* |  |
| Sinusoidal capillary lumen | 3 |
| Endothelial cell | 40 |
| Endothelial fenestration | 4 |
| Internal elastic lamina | 1 |
| *Portal vasculature* |  |
| Portal vein lumen | 1 |
| Portal endothelial cell | 8 |
| *Parenchyma, stroma, and extracellular matrix* |  |
| Hepatocyte | 50 |
| Stellate cell | 2 |
| Macrophage | 3 |
| Collagen bundles | 1 |

### 2.2 Automated segmentation enables construction of a labeled human liver volume map

To interpret the biological architecture of the reconstructed volume, we segmented the vasculature, cellular, and organelle structures across multiple spatial scales. Because these structures differ markedly in size, morphology, and density, we employed separate deep learning models optimized for each scale. For vascular and cellular structures, whose spatial coordinates were annotated in CATMAID, we used the Segment Anything Model 2 (SAM2) [15], a foundation model for prompt-based visual segmentation in images and videos (Fig. 2a). The coordinates from the annotated structure markers of each cellular structure (defined based on the middle of each structure instance) were used as point prompts for SAM2’s image mode, which generated pixel-wise labels corresponding to each structure, which were manually proofread. Using SAM2’s video mode, which uses a memory-attention mechanism that propagates object identity across frames, the initial label for each instance was extended through the volume toward both the beginning and end slices. This pipeline was applied to all annotated vascular and cell structures. Segmentation performance was evaluated against manual annotations using standard metrics including F1 score, precision, recall, intersection-over-union (IoU), and mean false distance [4, 8] (Table 2). The labels were then proofread to correct prediction error. The reconstructed labels were rendered in 3D to visualize the distribution of vascular and cellular structures within the periportal region (Fig. 2b, Supplementary Video 2).

**Table 2.** SAM2 performance metrics. Scores represent the mean of three tests against manually annotated ground truth. Higher values indicate better performance for F1 score, precision, recall, and IoU; lower values indicate better performance for MFD. IoU, Intersection over Union; MFD, Mean False Distance (pixels).

| Structure | F1 score | Precision | Recall | IoU | MFD |
| --- | --- | --- | --- | --- | --- |
| Bile duct lumen | 0.972 | 0.960 | 0.985 | 0.946 | 25.908 |
| Cholangiocyte | 0.949 | 0.985 | 0.917 | 0.903 | 186.889 |
| Endothelial cell | 0.862 | 0.947 | 0.791 | 0.758 | 168.338 |
| Hepatocyte | 0.979 | 0.993 | 0.965 | 0.959 | 222.484 |
| Portal vein lumen | 0.948 | 0.998 | 0.907 | 0.901 | 291.791 |
| Sinusoidal lumen | 0.953 | 0.930 | 0.977 | 0.909 | 43.928 |

**Fig. 2.**
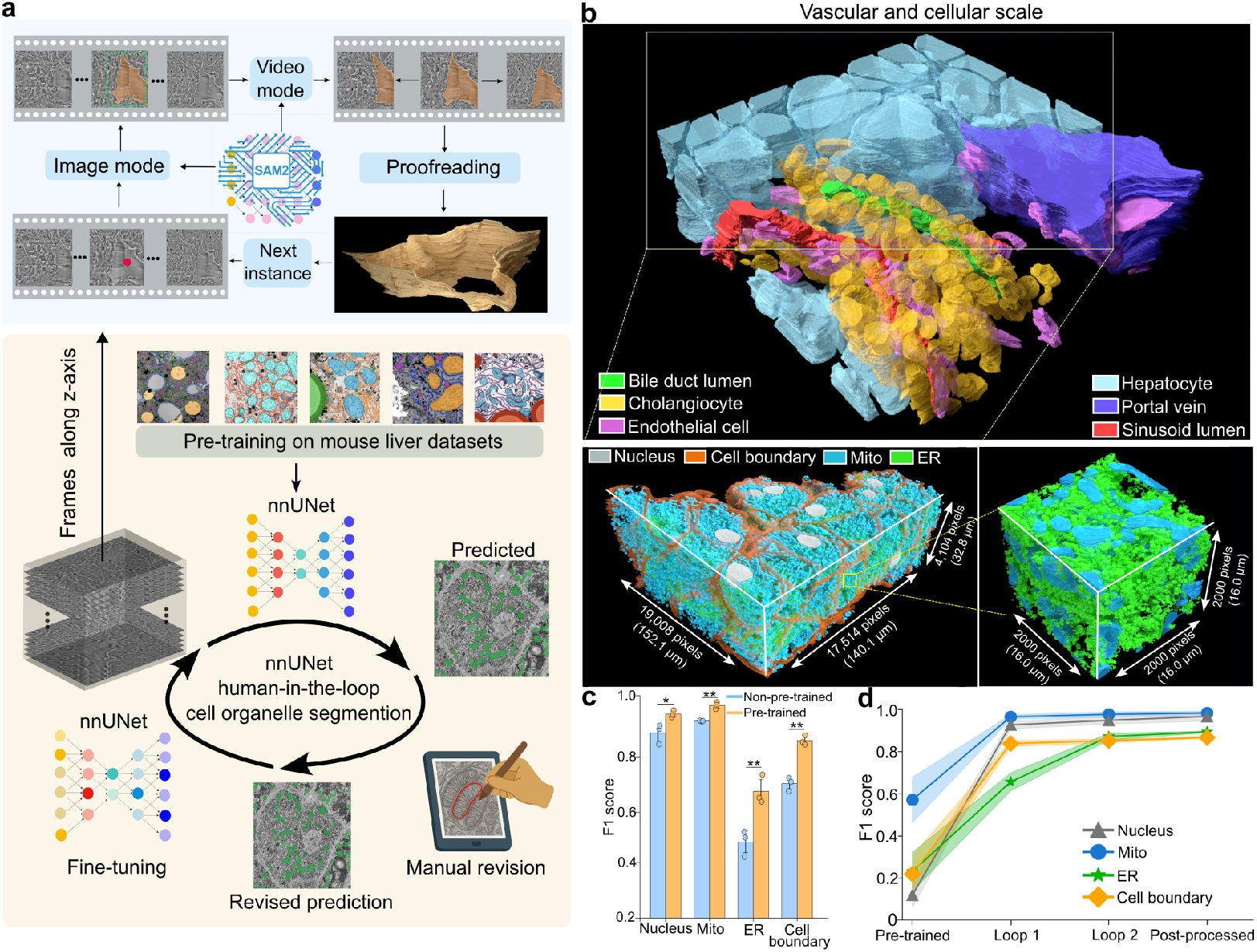
Automated deep learning segmentation of multiscale structures in the three-dimensional human liver volume. **a**, Workflow for multiscale structure annotation, from raw vEM images to labeled volumes. **b**, Three-dimensional rendering of the labeled volume showing multiscale segmented structures. **c**, Comparison of F1 scores between organelle segmentation models without pre-training versus models pre-trained on public vEM datasets, evaluated on held-out images (*n* = 3). **d**, Progressive improvement in segmentation accuracy (F1 score) across human-in-the-loop and post-processing steps. Statistical significance in **c** was determined by two-tailed paired t-test; \*\**P <* 0.01, \**P <* 0.05. Mito, mitochondria; ER, endoplasmic reticulum.

Unlike vascular and cellular structures that could be annotated with sparse spatial coordinates, subcellular organelles exhibited more regular morphologies but occurred in vast numbers throughout the volume, making coordinate-based prompting impractical. We therefore used nnU-Net V2 [16], a self-configuring semantic segmentation framework, to annotate organelles at the subcellular scale (Fig. 2a). Because pretraining improves segmentation accuracy on electron microscopy datasets [17, 18], we first pre-trained models on five publicly available mouse liver vEM datasets [4, 19]. We then implemented a human-in-the-loop training workflow to further improve labeling accuracy (Fig. 2a). In the first loop, a small subset of sections from the volume was manually annotated and used to train the models. In the subsequent loop, model predictions on different subsets were manually revised, and the corrected labels were used to retrain the model, followed by updated inference across the entire dataset. All manual annotations were performed using the Volume Annotation and Segmentation Tool (VAST) [20] (Supplementary Fig. 1g-h). Quantitative evaluation using F1 score, precision, recall, IoU, and mean false distance confirmed that pre-trained models after the first loop training significantly outperformed models without pre-training across four organelle classes (Fig. 2c, Table 3). We therefore continued with pre-trained models for subsequent loop training. After two loops, predictions underwent additional post-processing refinement using morphological operations (Table 4). Final performance substantially exceeded that of the models before human-in-the-loop training (Fig. 2d, Table 5). Organelle predictions were then reconstructed by stacking predicted section labels to produce the comprehensive multiscale human liver volume map (Fig. 2b, Supplementary Video 3).

**Table 3.** nnU-Net performance comparison between non-pre-trained and pre-trained models after one training loop. Scores represent the mean of three tests against manually annotated ground truth. Higher values indicate better performance for F1 score, precision, recall, and IoU; lower values indicate better performance for MFD. IoU, Intersection over Union; MFD, Mean False Distance (pixels); ER, endoplasmic reticulum.

| Structure | Model | F1 score | Precision | Recall | IoU | MFD |
| --- | --- | --- | --- | --- | --- | --- |
| Nucleus | Non-pre-trained | 0.865 | 0.871 | 0.859 | 0.762 | 2970.840 |
| Nucleus | Pre-trained | 0.926 | 0.942 | 0.910 | 0.862 | 2221.467 |
| Mitochondrion | Non-pre-trained | 0.908 | 0.915 | 0.901 | 0.832 | 2150.619 |
| Mitochondrion | Pre-trained | 0.964 | 0.980 | 0.949 | 0.931 | 1711.267 |
| ER | Non-pre-trained | 0.473 | 0.419 | 0.543 | 0.310 | 2257.592 |
| ER | Pre-trained | 0.655 | 0.610 | 0.709 | 0.487 | 1293.604 |
| Cell boundary | Non-pre-trained | 0.683 | 0.631 | 0.744 | 0.519 | 2841.694 |
| Cell boundary | Pre-trained | 0.837 | 0.788 | 0.893 | 0.720 | 1983.137 |

**Table 4.**
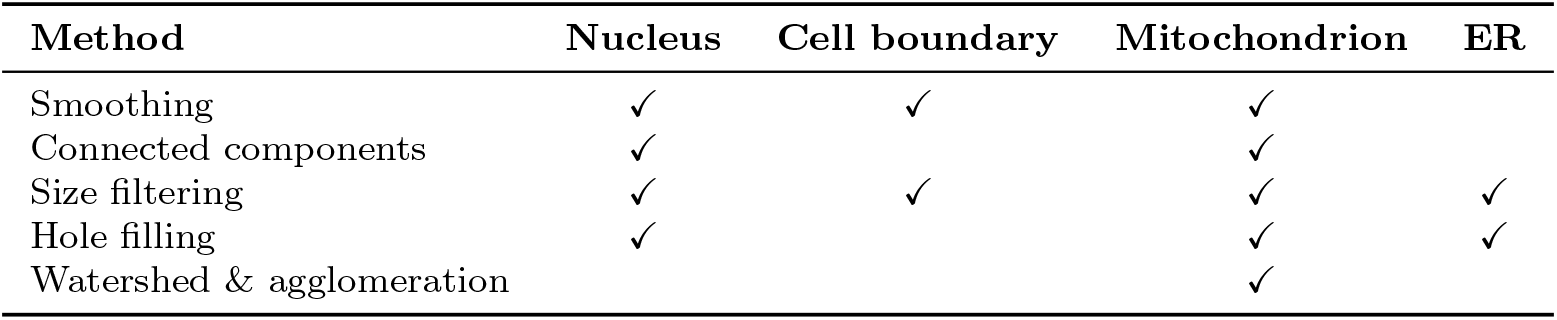
List of post-processing refinements used for organelles. ER: Endoplasmic Reticulum.

**Table 5.** nnU-Net performance metrics across iterative training loops. Scores represent the mean of three tests against manually annotated ground truth. Higher values indicate better performance for F1 score, precision, recall, and IoU; lower values indicate better performance for MFD. IoU, Intersection over Union; MFD, Mean False Distance (pixels); ER, endoplasmic reticulum.

| Label type | Stage | F1 score | Precision | Recall | IoU | MFD |
| --- | --- | --- | --- | --- | --- | --- |
| Nucleus | Pre-trained | 0.117 | 0.136 | 0.103 | 0.062 | 4256.676 |
| Nucleus | Loop 1 | 0.926 | 0.942 | 0.910 | 0.862 | 2221.467 |
| Nucleus | Loop 2 | 0.947 | 0.943 | 0.951 | 0.899 | 1398.563 |
| Nucleus | Post-processed | 0.966 | 0.951 | 0.981 | 0.934 | 844.835 |
| Mitochondrion | Pre-trained | 0.570 | 0.677 | 0.492 | 0.399 | 2927.612 |
| Mitochondrion | Loop 1 | 0.964 | 0.980 | 0.949 | 0.931 | 1711.267 |
| Mitochondrion | Loop 2 | 0.976 | 0.990 | 0.963 | 0.953 | 859.943 |
| Mitochondrion | Post-processed | 0.982 | 0.997 | 0.968 | 0.965 | 201.638 |
| ER | Pre-trained | 0.230 | 0.319 | 0.180 | 0.130 | 2075.234 |
| ER | Loop 1 | 0.655 | 0.610 | 0.709 | 0.487 | 1293.604 |
| ER | Loop 2 | 0.870 | 0.830 | 0.914 | 0.770 | 1125.926 |
| ER | Post-processed | 0.892 | 0.872 | 0.913 | 0.805 | 1099.278 |
| Cell boundary | Pre-trained | 0.217 | 0.308 | 0.168 | 0.122 | 3279.832 |
| Cell boundary | Loop 1 | 0.837 | 0.788 | 0.893 | 0.720 | 1983.137 |
| Cell boundary | Loop 2 | 0.851 | 0.826 | 0.878 | 0.741 | 1887.891 |
| Cell boundary | Post-processed | 0.866 | 0.843 | 0.890 | 0.764 | 1603.638 |

### 2.3 Cellular-scale analysis of the human liver map reveals lumen–cell relationships in bile ducts and sinusoidal capillaries

Bile ducts within the periportal zone transport bile and are lined by cholangiocytes, whose structural organization influences bile flow and composition [21]. Despite their importance in liver physiology and disease, detailed 3D analysis of bile duct architecture in human tissue remains limited. The bile duct comprises a central lumen surrounded by cholangiocytes (Fig. 3a). Across the serial EM slices, cholangiocytes were consistently observed encircling the lumen. Both the lumen diameter and the number of surrounding cholangiocytes increased progressively along the duct length (Fig. 3b). Quantitative analysis across the entire volume shows a positive correlation between lumen cross-sectional area and cholangiocyte number (Fig. 3c). To study lumen-cell contacts, we compared lumen cross-sectional area with average cholangiocyte cross-sectional area in individual slices (2D), and lumen cross-sectional area with average volume of nearby cholangiocytes across the duct (3D) (Fig. 3d, Supplementary Fig. 3a). Both approaches reveal a strong positive relationship between lumen size and cholangiocyte number and size, indicating coordination between the ductal lumen and its epithelial lining.

**Fig. 3.**
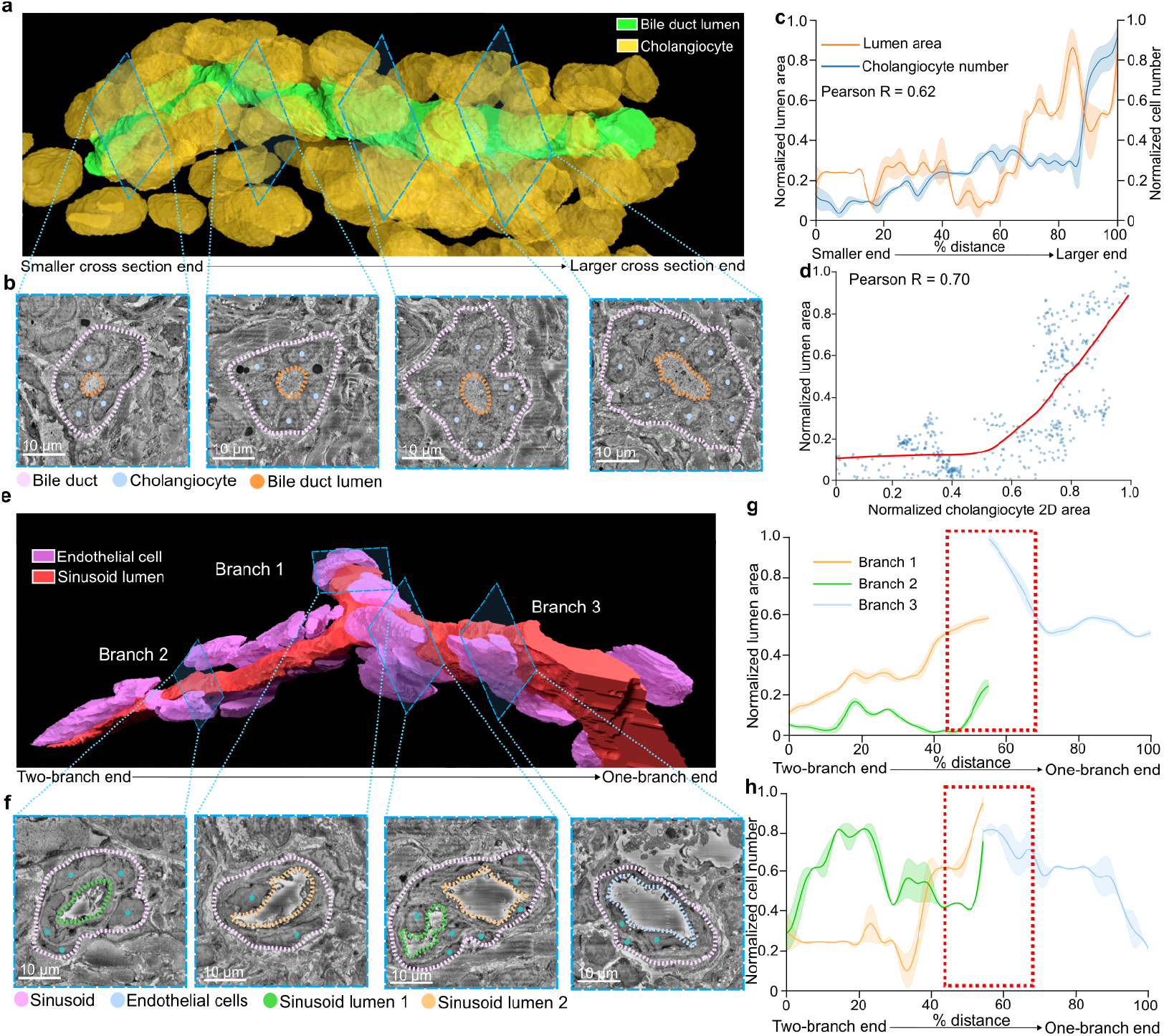
Lumen-cell relationships and branching morphology in biliary and sinusoidal vessels. **a**, Three-dimensional rendering of the bile duct lumen and surrounding cholangiocytes. Blue boxes indicate the positions of cross-sections shown in **b. b**, Representative serial EM cross-sections corresponding to the regions indicated in **a**, showing the bile duct lumen and adjacent cholangiocytes progressing from the smaller to larger cross-sectional area (left to right). **c**, Quantification of bile duct lumen area and surrounding cholangiocyte number across serial sections, progressing from the smaller to larger cross-sectional end (left to right). **d**, Correlation between bile duct lumen area and mean cholangiocyte area in two-dimensional slice-wise analysis. Red line represents locally weighted scatterplot smoothing (LOWESS) regression. **e**, Three-dimensional rendering of sinusoidal capillary lumen and surrounding endothelial cells. Blue boxes indicate the positions of cross-sections shown in **f. f**, Representative serial EM cross-sections corresponding to the regions indicated in **e**, showing the three sinusoidal branches (branch 1, branch 2, and branch 3) progressing from the two-branch end to the single-branch end (left to right). **g**, Lumen area of the three sinusoidal branches as a function of position along the vessel, measured from the two-branch end to the single-branch end. Red dashed box indicates the region near the branching point where the three branches converge. **h**, Endothelial cell number surrounding each sinusoidal branch as a function of position along the vessel. Red dashed box indicates the region near the branching point.

Beyond the bile duct system, sinusoidal capillaries constitute the primary microvasculature mediating blood-hepatocyte exchange. Sinusoidal branching patterns determine local blood flow dynamics and influence regional hepatocyte function [2]. We used the cellular-scale map capturing sinusoidal capillaries extending through the volume to investigate the change around a prominent branching point near the middle of the sinusoid where three distinct vascular segments converge (Fig. 3e, Supplementary Fig. 3b). Inspecting the 2D image slices resolved the spatial continuity of these segments, showing the gradual transition between two smaller branches and one larger branch (Fig. 3f). The two smaller segments (branch 1 and branch 2) display distinct morphologies: branch 1 exhibits a larger lumen surrounded by fewer endothelial cells (approximately two per cross-section), whereas branch 2 contains a smaller lumen enclosed by more endothelial cells (approximately five per section). The larger segment (branch 3) contains an intermediate number of endothelial cells (approximately three per section) (Fig. 3f). Three-dimensional quantitative tracing of the branches revealed that the lumen area of branch 1 exceeds that of branch 2, and that branch 3 approximates the combined lumen area of the two smaller branches, close to the branch point (Fig. 3g). Along the length of branch 3, however, lumen area progressively decreases, accompanied by reductions in both endothelial cell area and number (Fig. 3h, Supplementary Fig. 3c). To quantify the strength of endothelial-lumen contacts, we defined a contact ratio as the total cross-sectional area of endothelial cells divided by the lumen area, where higher values indicate greater endothelial coverage of the lumen volume. Branch 2 exhibits a significantly higher contact ratio than branch 1, while branch 3 displays an intermediate value closer to that of branch 1 (Supplementary Fig. 3d). This observation reveals asymmetric endothelial organization at the sinusoidal branching point where branch 2 exhibits high endothelial coverage whereas branch 1 and branch 3 display lower coverage.

### 2.4 Subcellular-scale analysis reveals mitochondrial morphological distribution

Beyond cellular and vascular structures, the subcellular-scale labeling of the multiscale volume revealed 35,790 complete mitochondria exhibiting diverse morphologies (Fig. 1f, Fig. 2b). To systematically characterize this morphological heterogeneity, we performed quantitative feature extraction and dimensionality reduction analysis. Shape-related morphological features of each mitochondrion were computed using PyRadiomics [22], which computes quantitative descriptors from 3D images (Table 6). Each mitochondrial instance was processed to generate a feature vector, which was subjected to principal component analysis (PCA) (Fig. 4a). PC1 accounts for the largest proportion of total variance (60.3%, Supplementary Fig. 4a). Mapping the extracted morphological features onto PCA space reveals that PC1 predominantly captures mitochondrial size, with higher PC1 values corresponding to increased surface area, mesh volume, and voxel volume (Fig. 4b, Supplementary Fig. 5). This trend was accompanied by concurrent changes in elongation (Fig. 4a-b, Supplementary Fig. 5). Visual inspection of representative mitochondria across the PC1 spectrum confirms that increasing PC1 values correspond to larger and more elongated morphologies (Fig. 4c). To examine the spatial distribution of morphologically distinct mitochondria within the volume, we selected representative instances from opposite tails of the PC1 distribution: low-PC1 instances from the left of the distribution peak and high-PC1 instances from the right (Supplementary Fig. 4b). Visualization of these examples in both two-dimensional cross-sections and three-dimensional subvolumes validated that PC1 reflects mitochondrial size and elongation (Fig. 4d-e). To characterize the spatial organization of morphologically distinct mitochondria at the whole-volume scale, we rendered the three-dimensional distribution of low-PC1 and high-PC1 mitochondria within individual hepatocytes. This analysis shows that both populations are distributed relatively homogeneously within each cell (Fig. 4f, Supplementary Video 4). We further quantified PC1 distributions within 30 hepatocytes whose cell bodies were largely contained within the imaged volume. Across all hepatocytes, the distributions consistently exhibit a leftward skew toward lower PC1 values with an extended tail toward higher values (Fig. 4g), mirroring the overall population distribution (Supplementary Fig. 4b). To assess whether mitochondrial morphology varies with hepatocyte position, we sorted hepatocytes by their distance to the portal vein and sinusoidal capillary; however, no distance-dependent gradient in PC1 distribution was observed (Fig. 4g, Supplementary Fig. 4c).

**Table 6.** Morphological features calculated by PyRadiomics.

| Feature name | Description |
| --- | --- |
| Mesh volume | Volume calculated from a 3D surface model of the region |
| Voxel volume | Volume estimated by counting the number of voxels within the region |
| Surface area | Total area of the region’s outer surface |
| Major axis length | Length of the longest axis of the best-fitting ellipsoid around the region |
| Minor axis length | Length of the second-longest axis of the best-fitting ellipsoid |
| Least axis length | Length of the shortest axis of the best-fitting ellipsoid |
| Max 3D diameter | Largest distance between any two points on the region surface |
| 2D diameter row | Largest distance between surface points in the sagittal plane |
| 2D diameter column | Largest distance between surface points in the coronal plane |
| 2D diameter slice | Largest distance between surface points in the axial plane |
| Sphericity | How closely the region resembles a sphere |
| Elongation | How stretched the region is along its two longest axes |
| Flatness | How flattened the region is, comparing the longest to the shortest axis |
| Surface volume ratio | Amount of surface area relative to the enclosed volume |

**Fig. 4.**
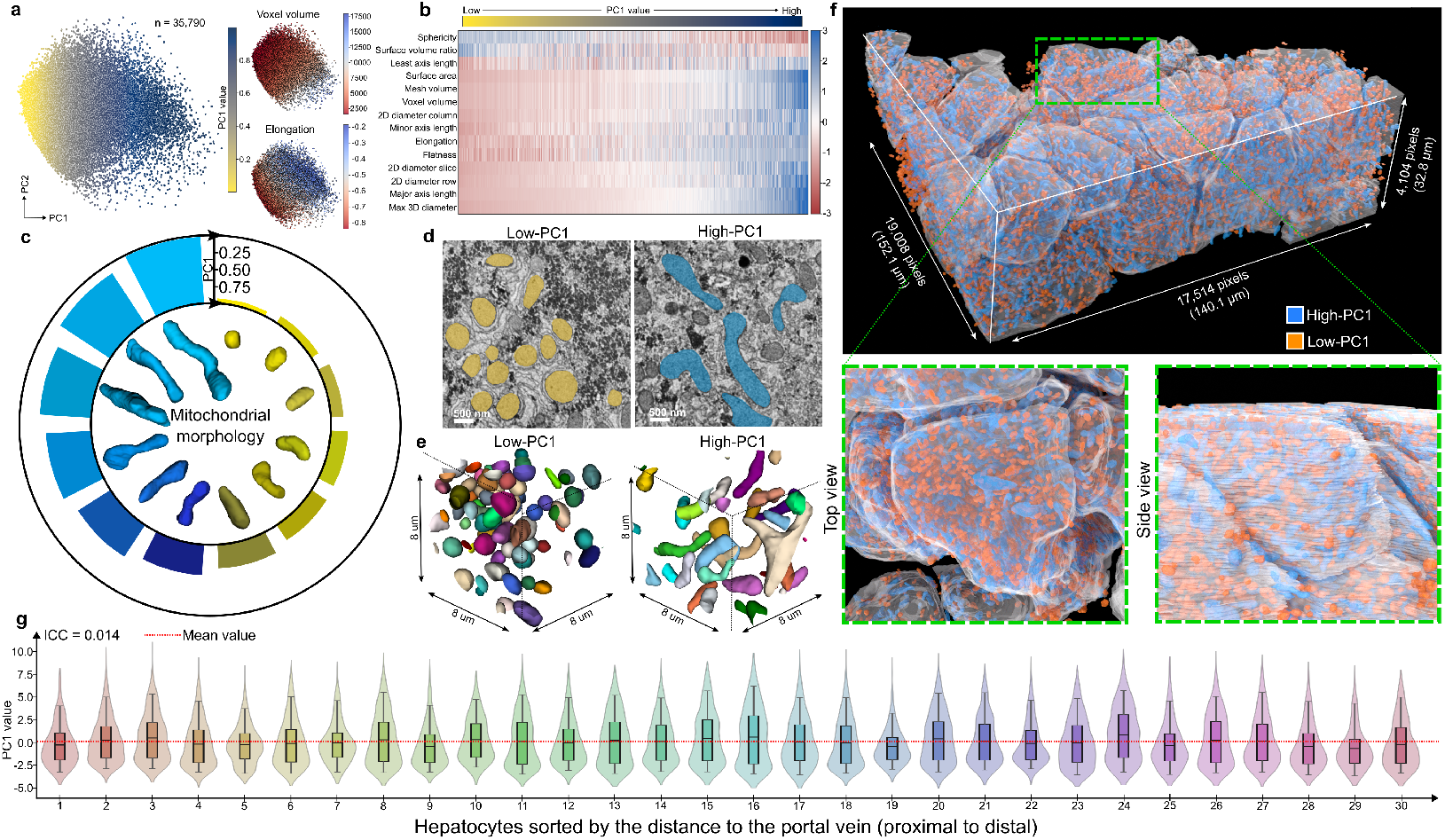
Mitochondrial morphological heterogeneity in the human liver volume. **a**, Left: principal component analysis (PCA) of 35,790 mitochondria based on extracted morphological features, with individual mitochondria colored by principal component 1 (PC1) values. Right: the same PCA projection colored by voxel volume (top) and elongation (bottom). **b**, Heatmap of morphological features across mitochondria sorted by PC1 value. Each column represents an individual mitochondrion; each row represents a morphological feature. **c**, Representative three-dimensional renderings of mitochondria spanning low to high PC1 values, with corresponding PC1 values indicated. **d**, Representative two-dimensional EM cross-section showing low-PC1 (orange) and high-PC1 (blue) mitochondria. **e**, Three-dimensional renderings of representative low-PC1 and high-PC1 mitochondrial instances. Individual mitochondria are shown in distinct colors. **f**, Three-dimensional visualization of low-PC1 and high-PC1 mitochondria across the entire volume. Hepatocyte boundaries are outlined in white. Green inset highlights a representative hepatocyte shown in top and side views. **g**, Violin plots showing PC1 distributions of mitochondria within 30 hepatocytes with relatively intact cell bodies. Hepatocytes are sorted by distance to the portal vein (left to right: proximal to distal). ICC, intraclass correlation coefficient.

We next examined PC2, which accounts for the second largest proportion of variance (18.8%, Supplementary Fig. 4a), to determine its relationship to mitochondrial morphology. Feature correlation analysis reveals strong associations between PC2 and both surface-to-volume ratio and least axis length (Supplementary Fig. 4d), metrics that describe the degree of local narrowing in mitochondrial shape (Table 6). To visualize morphological differences along the PC2 axis, we selected representative mitochondria from opposite tails of the PC2 distribution (Supplementary Fig. 4e). High-PC2 mitochondria display pronounced narrowing shapes compared to their low-PC2 counterparts (Supplementary Fig. 4f), consistent with the feature correlation analysis. Collectively, this PCA-based characterization reveals substantial heterogeneity in mitochondrial size, elongation, and narrowing morphology within the multiscale human liver volume, with conserved distributions across individual hepatocytes.

### 2.5 Mitochondria-ER contacts reveal mitochondrial dynamics

Given that mitochondria undergo continuous fission and fusion cycles [23], the observed morphological heterogeneity prompted us to investigate whether distinct morphological states reflect underlying mitochondrial dynamics. Because the ER has been implicated in regulating mitochondrial division and fusion events [23, 24], we analyzed the spatial relationships between mitochondria and ER in the human liver volume, beginning with comparisons across mitochondria of different size and elongation (PC1). To quantify mitochondria–ER contacts, we adopted a previously established distance-based method [25]. The ER surface was iteratively expanded at fixed distance intervals, and its intersection with the mitochondrial surface was measured at each step to quantify contact as a function of distance (Fig. 5a). Cryo-electron and transmission electron microscopy studies have shown that smooth mitochondria-ER contacts typically occur at 8-25 nm, whereas rough ER contacts can extend to 50-60 nm [25, 26]. Accordingly, we expanded the ER label at 32 nm (4 voxels) intervals up to 128 nm (16 voxels), at which distance most mitochondrial surfaces were covered by ER with minimal additional increase in contact beyond 96 nm (Fig. 5a). Two-dimensional and three-dimensional visualizations of representative high- and low-PC1 mitochondria reveal extensive ER surrounding both morphological classes (Fig. 5a-b). To determine whether mitochondria with distinct PC1 morphologies exhibit differential ER contacts, we compared the high- and low-PC1 populations defined in our PCA analysis. At short distances (0-32 nm), mitochondria with lower PC1 values exhibit significantly greater ER contact than those with higher PC1 values (Fig. 5c). This difference diminishes at intermediate distances (32-64 nm) and is almost absent at longer distances (64-128 nm), where both groups display comparable ER coverage. We next assessed whether this distance-dependent contact pattern is consistent across individual hepatocytes. In agreement with the population-level trend, low-PC1 mitochondria (smaller and more spherical) display significantly higher short-range (0-32 nm) ER contact than high-PC1 mitochondria (larger and more elongated) across most hepatocytes (Fig. 5d). At longer distances (96-128 nm), both populations exhibit comparable ER contact across hepatocytes. Together, these findings show that larger and elongated mitochondria maintain reduced short-range ER contact compared to smaller, spherical mitochondria, a pattern that is consistent across individual hepatocytes and reflects differential engagement with ER-mediated fission or fusion machinery.

**Fig. 5.**
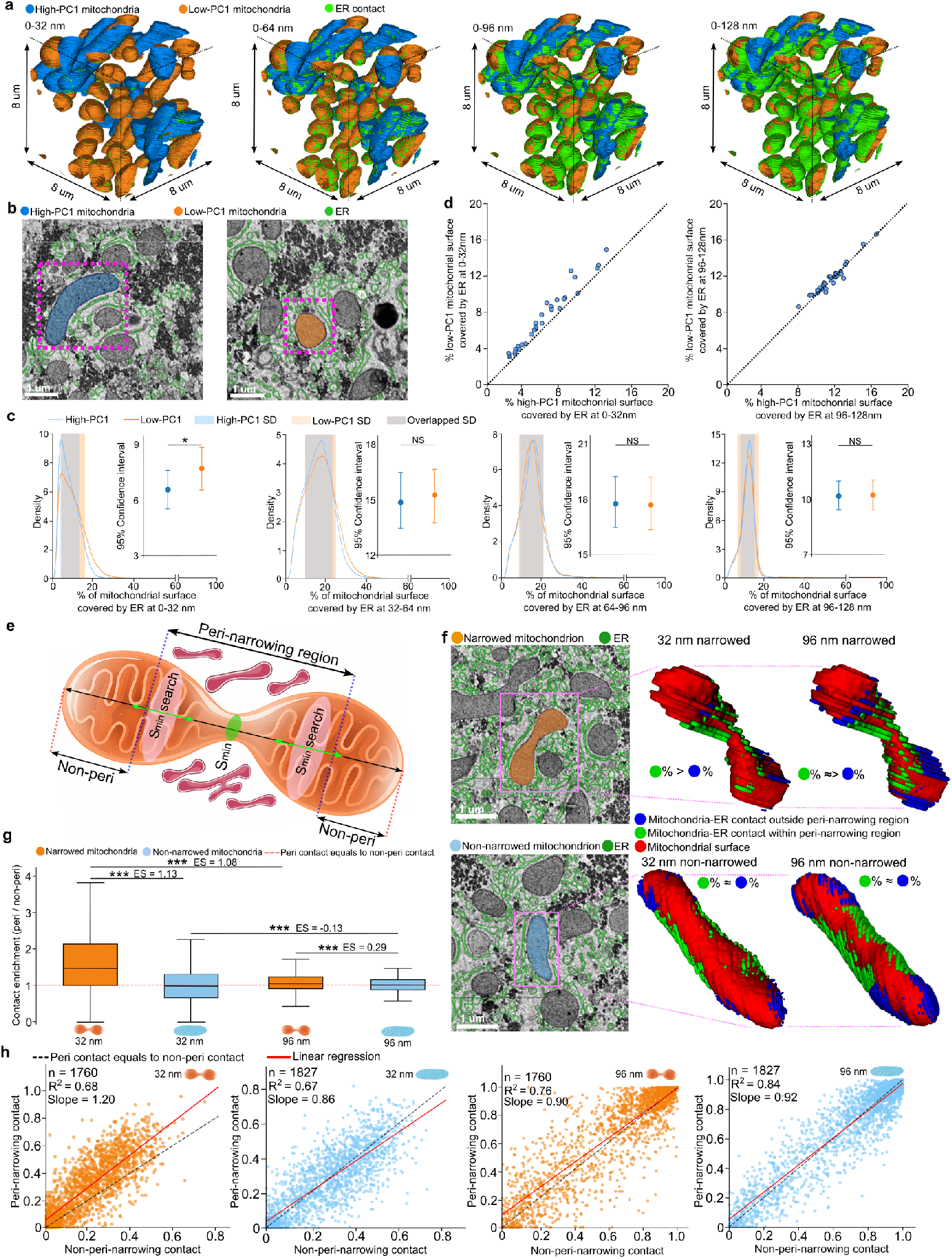
Mitochondria-endoplasmic reticulum (ER) contacts in the human liver volume. **a**, Three-dimensional visualization of mitochondria-ER contacts at increasing distance thresholds (0–128 nm in 32 nm intervals, left to right). **b**, Representative EM cross-sections showing ER surrounding high-PC1 (large and elongated; left) and low-PC1 (small and spherical; right) mitochondria. **c**, Distribution of the fraction of mitochondrial surface covered by ER, comparing high-PC1 and low-PC1 populations across distance thresholds (0-128 nm in 32 nm intervals, left to right). Kernel density estimates were computed from all individual mitochondria pooled across the volume (high-PC1, *n* = 12,351; low-PC1, *n* = 12,534). Inset: mean and 95% bootstrap confidence intervals computed from per-hepatocyte mitochondria-ER contact means. Statistical significance determined by two-tailed Mann-Whitney U test. **d**, Proportion of mitochondrial surface in contact with ER across 30 hepatocytes at short-range (0–32 nm; left) and long-range (96–128 nm; right) distances for high-PC1 and low-PC1 mitochondria. **e**, Schematic illustrating the identification of mitochondrial narrowing sites and the definition of peri-narrowing and non-peri-narrowing (Non-peri) regions used for quantifying local ER contact enrichment. **f**, Top row: representative narrowed mitochondrion shown in EM cross-section (left) and three-dimensional renderings illustrating ER contacts at 32 nm (middle) and 96 nm (right). Bottom row: representative non-narrowed mitochondrion with corresponding views. **g**, Box plots comparing ER contact enrichment between narrowed and non-narrowed mitochondria at 32 nm and 96 nm distances. Enrichment is defined as the ratio of ER-contacted surface area within the peri-narrowing region to that in the non-peri-narrowing region. Center line, median; box bounds, 25th–75th percentiles; whiskers, minimum and maximum values. Effect size calculated by pooled median absolute deviation; narrowed, *n* = 1, 760; non-narrowed, *n* = 1, 827. Statistical significance determined by two-tailed Mann-Whitney U test. **h**, Scatter plots showing the relationship between ER contact in peri-narrowing versus non-peri-narrowing regions for narrowed and non-narrowed mitochondria at 32 nm and 96 nm distances. Statistical significance indicated as \*\*\**P <* 0.001, \**P <* 0.05, NS not significant; PC, principal component; SD, standard deviation; ES, effect size.

Given the observed differences in mitochondria-ER contact level across morphologically distinct mitochondria, we next investigated whether these differences relate to mitochondrial dynamics. Previous studies have shown that elongated mitochondria often represent dynamic intermediates undergoing fission or fusion [7, 24], and that ER tubules mark mitochondrial division sites [6]. We therefore hypothesized that ER may be preferentially enriched at sites of mitochondrial narrowing in elongated mitochondria, reflecting active engagement with the fission or fusion machinery. To test this hypothesis, we focused on large, elongated mitochondria (high PC1) and stratified them by narrowing morphology using PC2, which captures the degree of local narrowing (Supplementary Fig. 6a). High-PC2 mitochondria, characterized by prominent narrowing sites, were designated as narrowed mitochondria, while low-PC2 mitochondria lacking such features served as non-narrowed controls. For each narrowed mitochondrion, we identified the narrowing site as the position of minimal cross-sectional area along the major axis. The surrounding 50% of the mitochondrial surface centered on this site was defined as the peri-narrowing region, with the remainder designated as the non-peri-narrowing region (Fig. 5e). Mitochondria lacking a clearly defined local minimum in cross-sectional area were excluded from subsequent analyses. Visual inspection reveals distinct patterns of ER distribution between the two populations. For narrowed mitochondria, ER coverage at 32 nm is notably higher in peri-narrowing regions compared to non-peri-narrowing regions, whereas this difference is diminished at 96 nm (Fig. 5f, Supplementary Fig. 6b). In contrast, non-narrowed mitochondria exhibit no apparent spatial preference for ER contact at either distance.

To quantify these observations, we calculated the ER contact density (ER-contact surface area divided by total mitochondrial surface area) separately for peri-narrowing and non-peri-narrowing regions (Supplementary Fig. 6c). Narrowed mitochondria exhibit significantly higher ER contact density in peri-narrowing regions, with a larger effect size at 32 nm than at 96 nm. Non-narrowed mitochondria show no significant regional difference in ER contact density. We further computed an enrichment ratio, defined as the ER contact density in the peri-narrowing region divided by that in the non-peri-narrowing region. At 32 nm, narrowed mitochondria predominantly display enrichment ratios greater than 1, indicating preferential ER association near narrowing sites (Fig. 5g-h). Non-narrowed mitochondria, by contrast, exhibit enrichment ratios clustered around 1. At 96 nm, enrichment ratios for narrowed mitochondria remain modestly above 1, while those for non-narrowed mitochondria approximate 1, indicating no preferential ER localization. Together, these findings demonstrate that mitochondria with narrowing sites exhibit preferential short-range ER contact at these sites, consistent with ER involvement in mitochondrial remodeling. This spatial relationship provides ultrastructural evidence of mitochondrial dynamics that reflect fission or fusion intermediates preserved within the human liver tissue volume.

## 3 Discussion

We present a multiscale, multicellular vEM image of a human liver periportal region, establishing a nanoscale reference atlas spanning tissue, vascular, cellular, and subcellular organization. To achieve this, we developed an integrated imaging pipeline that accommodates multiscale region selection and acquisition across spatial scales. Using deep learning-based segmentation, we generated a comprehensive three-dimensional annotation encompassing vasculature, cells, and organelles. This multilevel reconstruction enables quantitative analyses across spatial scales, revealing structural heterogeneity at multiple levels of organization: vascular architecture including bile duct and sinusoidal branching patterns, cellular-scale features such as endothelial coverage and cholangiocyte distribution, and organelle-level characteristics including mitochondrial morphological diversity and spatially enriched mitochondria-ER contacts at sites of mitochondrial narrowing, consistent with signatures of mitochondrial dynamics.

The observed correlation between bile duct lumen size and cholangiocyte number and size aligns with mechanobiological models of ductal morphogenesis. Previous studies have shown that bile duct lumen formation depends on coordinated cholangiocyte proliferation, osmotic forces from biliary secretions, and timed differentiation, with luminal expansion emerging from these processes [21]. Our finding that lumen enlargement is accompanied by an increase in surrounding cholangiocytes is consistent with these mechanisms. This relationship also provides a framework for understanding pathologies such as vanishing bile duct syndrome, where ductopenia (the loss or absence of bile ducts per portal tract) reflects progressive cholangiocyte depletion and precedes cholestatic disease progression [27, 28]. The strong structural correlation between lumen size and cholangiocyte abundance in our dataset thus mirrors clinical observations of bile duct loss in cholangiopathies, suggesting a potential morphological basis for early diagnostic indicators.

The asymmetric sinusoidal branching observed in the labeled volume, where a thinner branch with extensive endothelial-lumen contact converges with a wider branch exhibiting reduced endothelial coverage, suggests localized adaptation to mechanical stress. Liver sinusoidal endothelial cells are highly mechanosensitive, modulating fenestration size and number in response to flow dynamics to regulate material exchange between sinusoidal blood and hepatocytes [29]. Our ultrastructural observations captured fenestrated endothelium lining the sinusoidal capillary (Supplementary Fig. 2a) and stellate cells residing in the perisinusoidal space between hepatocytes and endothelium (Supplementary Fig. 2b), suggesting active exchange function in the periportal region. Reduced endothelial interaction in the wider branch may optimize hemodynamic efficiency while minimizing shear-induced damage. However, disturbed shear stress can drive endothelial dysfunction and capillarization, key contributors to microvascular remodeling and fibrosis [30]. Consistent with prior observations that periportal endothelial cells display a lower abundance of fenestration with larger diameters, favoring selective filtration and oxygen delivery, whereas pericentral endothelial cells exhibit more numerous but smaller fenestration that optimize rapid exchange and waste clearance [31], we propose that the thinner, high-interface branch represents a metabolically active exchange zone, while the wider, low-interface branch functions as a mechanically stable transport conduit. Disruption of this balance between exchange efficiency and mechanical stability may contribute to microvascular dysfunction, as impaired substance exchange resulting from fenestration loss is a recognized early event in liver disease that precedes hepatic stellate cell activation and fibrogenesis [32, 33]. However, validation of this hypothesis requires comparative analysis with diseased tissue.

Quantitative analysis of 35,790 mitochondria and their ER contacts provides ultra-structural evidence consistent with ER-mediated regulation of mitochondrial dynamics in human liver tissue. Previous studies in cell culture and animal models have demonstrated that ER tubules mark mitochondrial division sites and remain closely associated during fission and fusion events, with smooth ER specifically implicated at these sites [6, 7, 23, 25]. Although not all elongated mitochondria in our dataset necessarily represent active fission or fusion intermediates captured in a static snapshot, statistical analysis across this large population revealed consistent enrichment of ER contacts at narrowing sites, aligning with these prior observations. This correlation suggests that a subset of the morphological heterogeneity observed here reflects ongoing mitochondrial dynamics preserved as ultrastructural signatures in human liver tissue.

The imaging resolution and contrast in our dataset precluded reliable identification of ribosomes to distinguish smooth from rough ER. Nevertheless, the preferential ER enrichment at mitochondrial narrowing sites falls within the distance range reported for smooth ER contacts [25, 26]. Future studies employing higher-resolution imaging or correlative approaches will be necessary to definitively characterize ER subtype involvement in human tissue. Additionally, we did not detect zonation-dependent differences in mitochondrial morphology across hepatocytes, potentially because our dataset was limited to the periportal region. Extending this approach to liver tissue volumes spanning the porto-central axis will determine whether mitochondrial morphological gradients exist in human liver.

Collectively, this work provides an unprecedented multiscale reconstruction of human liver architecture, establishing both a structural reference for the healthy peri-portal region and a computational framework for quantitative organelle analysis. The imaging and analysis pipeline developed here will facilitate comparative studies across hepatic zones and disease states, enabling investigation of how tissue, vascular, cellular, and organelle organization becomes altered in liver pathology. Such multiscale atlases, integrating automated segmentation with quantitative morphological analysis, represent a powerful approach for bridging ultrastructural observations to functional and pathophysiological insights.

## 4 Methods

### 4.1 Human liver tissue acquisition

Human liver tissue was obtained from a neurologically deceased donor deemed acceptable for liver transplantation. Sample collection was performed under institutional ethics approval (University Health Network REB# 14-7425-AE) with de-identified donor metadata recorded and stored securely, as previously described [34].

Donor livers were flushed *in situ* with cold histidine-tryptophan-ketoglutarate (HTK) preservation solution. The caudate lobe was resected and maintained in HTK solution on ice. In the laboratory, the lobe was cannulated using 1.2–2.0 mm irrigation cannulae inserted into exposed vessels on the cut surface and gently perfused at 4°C with HEPES-buffered saline (HBS) containing EGTA (approximately 10 mL min^−1^ per cannula) to clear residual blood.

For fixation, tissue was transferred to a chilled Petri dish, trimmed into approximately 1 mm^3^ cubes using a sterile single-edge razor blade, and immediately immersed in aldehyde fixative (4% formaldehyde and 2.5% glutaraldehyde in 0.1 M cacodylate buffer, pH 7.4) for 12–24 h at 4°C. After fixation, samples were rinsed thoroughly in 0.1 M cacodylate buffer (3 *×* 10 min).

### 4.2 Tissue processing for volume electron microscopy

Aldehyde-fixed tissues were processed using a modified OTO (osmium-thiocarbohydrazide-osmium) protocol. Samples were first post-fixed in 2% osmium tetroxide (OsO_4_) with 1% potassium ferrocyanide (K_4_[Fe(CN)_6_]) in 0.1 M cacodylate buffer for 90 min at room temperature. After washing with distilled water, samples were stained with 1% thiocarbohydrazide (TCH) for 30 min, rinsed, and stained with 1% osmium tetroxide for 1 h. Tissue was then washed and dehydrated through a graded ethanol series (50%, 70%, 90%, 100%; 20 min each step) and equilibrated in propylene oxide. Infiltration proceeded in a 1:1 mixture of propylene oxide and Spurr resin for 2 h, followed by 100% Spurr resin overnight at room temperature. The following day, samples were transferred to fresh Spurr resin for an additional 2 h, embedded in silicone molds, and polymerized at 70°C for 24 h.

### 4.3 X-ray micro-computed tomography

X-ray micro-computed tomography was performed on a Zeiss Xradia Versa 510 X-ray microscope to assess sample integrity and identify regions of interest prior to electron microscopy. The resin-embedded specimen was imaged at 80 kV with 360° sample rotation to acquire projection images for three-dimensional reconstruction.

### 4.4 Immunohistochemistry

A portion of the same liver specimen was processed for immunohistochemistry to confirm hepatic zonation. Tissue was cut into 4 mm *×* 4 mm *×* 4 mm blocks and fixed in 10% neutral buffered formalin. Formalin-fixed, paraffin-embedded tissue was sectioned at 5–7 µm thickness. Immunohistochemical staining was performed on sequential sections by the Toronto Pathology Research Program (Toronto General Hospital) using standard protocols. Slides were pretreated with low-temperature target retrieval solution (pH 9) for antigen retrieval. Primary antibody against CYP2A6, a periportal marker enzyme, was applied, followed by detection using donkey anti-rabbit secondary antibody conjugated to horseradish peroxidase. Slides were digitized using a ScanScope AT2 scanner (Leica Biosystems) at *×* 40 magnification at the Advanced Optical Microscopy Facility, Toronto. Periportal and pericentral regions were confirmed by a board-certified liver pathologist (C.T.).

### 4.5 Serial block-face scanning electron microscopy

Imaging was performed on a Zeiss Gemini 2 SEM equipped with a Gatan 3View 2XP ultramicrotome. Serial sectioning and image acquisition were automated using 3View control software. Images were acquired at an accelerating voltage of 2.5 kV with a 30 µm objective aperture, the electron gun in analytical mode, and high-current beam settings. Backscattered electron signal was detected using the in-column detector to enhance heavy-metal contrast. Focal charge compensation with nitrogen gas was applied during imaging to mitigate charging artifacts. For each section, the dwell time was 3 µs per pixel with 8 nm pixel size. The sectioning thickness (z-step) was 55 nm throughout the acquisition. Autofocus and stigmator correction routines were engaged at regular intervals, and drift correction was applied between cycles as needed. The resulting dataset comprised a contiguous three-dimensional volume with 8 nm in-plane pixel size and 55 nm axial spacing.

### 4.6 EM image stitching and alignment

Stitching and alignment were performed in Python using Google’s SOFIMA algorithm (https://github.com/google-research/sofima) [13]. For stitching, we processed 5 *×* 5 tile arrays for each section of the serial block-face scanning electron microscopy dataset. Coarse tile position optimization was first applied to roughly define relative tile positions by solving a constrained optimization problem. The obtained relative positions from the coarse optimization step were then used to compute flow fields between pairs of neighboring tiles to achieve fine tile position optimization. For mesh solving, each tile was modeled as a spring mesh, and the equilibrium state was calculated to balance two competing forces: the spring elastic force, which preserves the original tile geometry, and a force associated with the calculated flow field, which ensures connectivity with neighboring tiles. Based on the solved mesh, the optimally positioned tiles were rendered to generate the final stitched section.

For alignment, flow fields were computed between each stitched section and its adjacent sections. Both full-resolution and 2*×* downsampled flow fields were calculated, as the lower-resolution flow field can compensate for regions where the full-resolution flow field estimation is not reliable. Uncertain flow estimates were filtered out. For mesh optimization, an elastic mesh optimizer was used to balance the estimated flow field with the preservation of original biological structure geometry. This optimization was applied sequentially, section by section. Finally, the sections were rendered and the volume was reconstructed by stacking the aligned rendered sections.

### 4.7 Liver structure annotation

To facilitate visualization and annotation, the vEM dataset was imported into a CAT-MAID [14] server hosted by the Zhen lab. Structures in each section were manually annotated and validated by a board-certified liver pathologist (C.T.). Annotations included different cell types and vascular structures. Point markers were placed at the center of each annotated instance to record spatial coordinates. Sinusoidal endothelial cells present unique annotation challenges due to their discontinuous architecture, characterized by gaps between cells and fenestrations [35]. Therefore, endothelial cells were identified based on their nuclear morphology. Cytoplasmic fragments without visible nuclei, which represent the discontinuous nature of sinusoidal architecture, were not independently quantified as separate cellular entities. All annotated structure types and their coordinates are available through the Zenodo link provided in the Data Availability section.

### 4.8 Manual segmentation of subcellular structures

Following pathologist validation, manual segmentation was performed on selected sections. Target sections were imported into VAST [20], a tool for labeling large image stacks. We used the pen tool to manually annotate nuclei, cell boundaries, mitochondria, and ER within hepatocyte regions. The segmentation labels were then exported from VAST for downstream deep learning model training.

### 4.9 Vascular and cellular scale segmentation

Vascular structures (sinusoidal capillary lumen, bile duct lumen, and portal vein lumen) and cellular structures (endothelial cells, cholangiocytes, and hepatocytes) were segmented using SAM2 [15], a foundation model for promptable segmentation of images and videos. Annotated point prompts were placed in CATMAID on each target structure, positioned at the middle cross-section of objects spanning the z-axis. In image mode, the model predicted the label of the current object, which was then proofread. Based on this label, a bounding box was defined by recording the left-most, topmost, rightmost, and bottommost coordinates. These bounding boxes were input to the model in video mode to propagate the predicted labels from the initial section to both the start and end of the instance across the volume, achieving complete three-dimensional segmentation of each object. To evaluate segmentation performance, three randomly selected cross-sections of each object were manually annotated and compared with the corresponding model predictions. All labels underwent final proof-reading. Certain cellular structures presented segmentation challenges due to their inherent morphology. Sinusoidal endothelial cells possess extremely thin cytoplasmic extensions and fenestrated regions where cell boundaries are not clearly delineated at the imaging resolution. Similarly, cholangiocytes exhibit irregular apical surfaces with microvillar projections that create ambiguous cell boundaries. For these cell types, conservative segmentation was applied, annotating only regions with clearly identifiable cell structures to avoid over-annotation. Consequently, the segmented cell volumes may underestimate true cell size in regions of thin cytoplasm or complex membrane topology.

### 4.10 Subcellular scale segmentation

Nuclei, mitochondria, ER, and hepatocyte boundaries were automatically segmented using nnU-Net [16], a self-configuring U-Net model. The model processes individual 2D image slices as input and predicts pixel-wise segmentation. All hyperparameters and model configurations follow the nnU-Net self-configured settings. Each segmentation class was trained with a separate model. Models of nucleus, mitochondrion, and ER were pre-trained on five mouse liver datasets from OpenOrganelle [19] (dataset IDs: jrc mus-liver, jrc mus-liver-4, jrc mus-liver-5, jrc mus-liver-6, and jrc mus-liver-7). Model of cell boundary was pre-trained on one mouse liver dataset from OpenOrganelle (dataset ID: jrc mus-liver) After pre-training, the neural network encoder was frozen and the model was further trained on our manually annotated human liver images. Human-in-the-loop refinement was performed by manually correcting the predicted segmentation on images not included in the current training set. These revised segmentations and their corresponding images were then added to the training data for subsequent iterations. This process was repeated for two loops. Finally, the predicted segmentations were post-processed using a series of morphological operations Supplementary (Table 4). The segmentations were smoothed using a Gaussian filter (scipy.ndimage.gaussian filter) to reduce boundary irregularities. Connected component analysis (skimage.measure.label) was performed to identify individual objects. Size filtering was used to remove small spurious fragments (skimage.morphology.remove small objects). Internal holes within segmented structures were filled (scipy.ndimage.binary fill holes) to ensure topological consistency. And watershed-based agglomeration was applied to merge over-segmented regions (skimage.segmentation.watershed). For segmentation evaluation, we randomly selected three fully manually segmented sections that were excluded from training. All classes were evaluated by comparing manual and automated segmentation on these test images. Model performance was assessed at each stage (pre-trained models, each loop training, and final segmentation) using F1 score (Dice coefficient), intersection-over-union (IoU), precision, recall, and mean false distance based on the Hausdorff distance.

### 4.11 vEM volume and label visualization

The vEM volume and segmented labels were visualized using Blender [36] (version 4.3.2) with the Microscopy Nodes add-on for microscopy data visualization. The vEM volume was converted to TIFF format in Python and imported into Blender using the volume option in Microscopy Nodes. Segmentation labels were separately converted to binary TIFF images and imported using Microscopy Nodes with surface and emission rendering. Videos were generated by setting up animations in Blender. Subvolumes of mitochondrial labels and 3D visualizations of mitochondria–ER contacts (Fig. 4 and Fig. 5d) were created using ITK-SNAP [37], an interactive tool for segmentation visualization.

### 4.12 Bile duct lumen and cholangiocyte characterization

Morphometric measurements of bile duct lumens and cholangiocytes were obtained using the skimage.measure module. To quantify cholangiocyte distribution, cells were identified within a distance threshold of 1500 voxels from the bile duct lumen surface. For each cross-section of the bile duct lumen, two complementary analyses were performed: quantification of cholangiocytes within the same cross-sectional plane, and quantification of intact three-dimensional cholangiocytes located within adjacent cross-sections. This dual approach enabled assessment of both local cellular organization and the overall spatial relationship between cholangiocytes and the bile duct lumen.

### 4.13 Sinusoidal capillary lumen and endothelial cell characterization

Morphometric measurements of sinusoidal capillary lumens and associated endothelial cells were obtained using the skimage.measure module. Endothelial cells were identified within a distance threshold of 1500 voxels from the lumen surface. The sinusoidal branching point was identified near slice index 281, where two branches converge into a single vessel. For each of the three branches, we quantified both the lumen cross-sectional area and the number and size of surrounding endothelial cells. This branch-specific analysis enabled comparison of endothelial coverage across different segments of the sinusoidal network.

### 4.14 Mitochondrial morphological feature extraction

Following mitochondrial segmentation, connected component analysis was applied to assign a unique identifier to each mitochondrion, yielding 37,920 objects. Mitochondria touching any volume boundary were excluded to ensure only complete, non-truncated ones were analyzed, leaving 35,790 mitochondria for downstream analysis. Morphological features were extracted using PyRadiomics [22]. The RadiomicsShape three-dimensional descriptors were used to characterize mitochondrial size and shape, including: Elongation, Flatness, LeastAxisLength, MajorAxisLength, Maximum2DDiameterColumn, Maximum2DDiameterRow, Maximum2DDiameterSlice, Maximum3DDiameter, MeshVolume, MinorAxisLength, Sphericity, SurfaceArea, SurfaceVolumeRatio, and VoxelVolume. Feature descriptions are provided in Table 6.

### 4.15 Principal component analysis for mitochondrial morphology characterization

To characterize mitochondrial morphology, the 14 extracted features were standardized using z-score normalization (StandardScaler) prior to principal component analysis (PCA). PCA was performed using the scikit-learn Python package. The distributions of the first and second principal components (PC1 and PC2) across all mitochondria within the volume were fit using a skew-normal distribution (scipy.stats.skewnorm). PC1 captured morphological variation along a spectrum from compact to elongated mitochondria, while PC2 distinguished mitochondria with and without narrowing regions. For visualization, individual mitochondria were classified based on their PC scores. High-PC2 mitochondria were defined as those with PC2 > 1, representing mitochondria with prominent narrowing sites, while low-PC2 mitochondria were defined as those with PC2 < -1, representing mitochondria lacking such features. The segmentation masks of representative mitochondria from each group were extracted and rendered in ITK-SNAP for three-dimensional visualization.

### 4.16 Mitochondria-ER contact characterization

Mitochondria-ER contacts were characterized by measuring the distance between mitochondrial surfaces and ER. Following a previously established method [25], the mitochondrial outer surface was generated by eroding the mitochondrial volume by one voxel and subtracting the eroded volume from the original volume. ER labels were then iteratively dilated based on the desired distance intervals. The number of overlapping voxels between the dilated ER and mitochondrial surface was counted as the interacting surface. The percentage of mitochondrial surface covered by ER was calculated by dividing the interacting surface area by the total mitochondrial surface area. To avoid multiple counting, voxels counted in a given dilation step were excluded from subsequent larger dilation steps.

### 4.17 Mitochondrial dynamics in relation to narrowing sites

Mitochondria were filtered based on their morphological characteristics as captured by principal component analysis (PCA). Only mitochondria with PC1 values greater than 1 were included in the analysis. These were further subdivided into narrowing mitochondria (PC2 > 1) and non-narrowing mitochondria (PC2 < -1). To identify narrowing sites, the major axis of each mitochondrion was determined by performing PCA on the 3D coordinates of its constituent voxels. Cross-sectional areas were calculated perpendicular to this axis at regular intervals. A narrowing site was defined as a position exhibiting a local minimum in cross-sectional area, excluding the terminal regions where cross-sectional area naturally decreases toward the mitochondrial endpoints. The local minimum was further validated by requiring it to be less than 80% of the median cross-sectional area along the entire axis. When multiple local minima were present, the position with the smallest cross-sectional area was selected as the narrowing site. For narrowing mitochondria, the peri-narrowing region was defined as the 50% of the mitochondrial surface centered on the narrowing site. For non-narrowing mitochondria lacking an identifiable local minimum in cross-sectional area, the peri-narrowing region was defined as the middle 50% of the surface (corresponding to 25–75% along the major axis). In both cases, the remaining surface constituted the non-peri-narrowing region. Contact enrichment was calculated as the ratio of mitochondria–ER contact density in the peri-narrowing region to that in the non-peri-narrowing region, where contact density was defined as the percentage of mitochondrial surface in contact with ER. Values greater than 1 indicate preferential ER contact at or near narrowing sites.

### 4.18 Statistics and reproducibility

Statistical analyses were carried out using Python. Differences between groups were determined using unpaired t-test and p-values are indicated in the figure legends. Numbers of biological replicates are indicated in the figure legends. All experiments were repeated in at least in duplicate. No statistical methods were used to predetermine sample size.

## Supplementary material

Supplementary material includes figures and videos.

## 5 Data availability

Raw FIB-SEM images and their segmentation have been deposited to the EMPIAR [38] database, available using accession code: EMPIAR-13356. The manually labeled structures and the model trained weights are available at Zenodo. Supplementary videos are available at Google Drive or Zenodo.

## 6 Code availability

The source code generated and analyzed in this study is available at GitHub.

## Acknowledgements

All authors thank the Digital Research Alliance of Canada for high-performance computing access.

## Author contributions

C.X., R.X., and G.D.B. conceptualized the study and designed the project. C.X. performed automated segmentation and computational analysis, prepared figures, and wrote the manuscript. B.M., A.D., I.S.C., and M.Z. performed, supervised, and executed volume electron microscopy data acquisition. J.M. advised on the deep learning pipeline. C.T., Y.L., and F.A. performed manual image annotation. I.D.M. and S.M. performed tissue sample acquisition. G.D.B. conceived and supervised the project, secured funding, interpreted results, and revised the manuscript.

## Supplementary information

The supplementary material is available in the PDF file attached.

## References

[1] Arias, I. M. et al. (eds) The Liver: Biology and Pathobiology (John Wiley & Sons, Hoboken, NJ, 2020).

[2] Segovia-Miranda, F. et al. Three-dimensional spatially resolved geometrical and functional models of human liver tissue reveal new aspects of NAFLD progression. Nature Medicine 25, 1885–1893 (2019).

[3] Morales-Navarrete, H. et al. Liquid-crystal organization of liver tissue. Elife 8, e44860 (2019).

[4] Parlakgül, G. et al. Regulation of liver subcellular architecture controls metabolic homeostasis. Nature 603, 736–742 (2022).

[5] Trimm, E. & Red-Horse, K. Vascular endothelial cell development and diversity. Nature Reviews Cardiology 20, 197–210 (2023).

[6] Friedman, J. R. et al. ER tubules mark sites of mitochondrial division. Science 334, 358–362 (2011).

[7] Kleele, T. et al. Distinct fission signatures predict mitochondrial degradation or biogenesis. Nature 593, 435–439 (2021).

[8] Han, M. et al. Spatial mapping of mitochondrial networks and bioenergetics in lung cancer. Nature 615, 712–719 (2023).

[9] Takahashi, K. et al. An analysis modality for vascular structures combining tissue-clearing technology and topological data analysis. Nature Communications 13, 5239 (2022).

[10] Karschau, J. et al. Resilience of three-dimensional sinusoidal networks in liver tissue. PLOS Computational Biology 16, e1007965 (2020).

[11] Kang, S. W. S. et al. A spatial map of hepatic mitochondria uncovers functional heterogeneity shaped by nutrient-sensing signaling. Nature Communications 15, 1799 (2024).

[12] Halpern, K. B. et al. Single-cell spatial reconstruction reveals global division of labour in the mammalian liver. Nature 542, 352–356 (2017).

[13] Januszewski, M., Blakely, T. & Lueckmann, J.-M. Sofima (scalable optical flow-based image montaging and alignment). Zenodo10 5281 (2023).

[14] Saalfeld, S., Cardona, A., Hartenstein, V. & Tomančák, P. Catmaid: collaborative annotation toolkit for massive amounts of image data. Bioinformatics 25, 1984–1986 (2009).

[15] Ravi, N. et al. Sam 2: Segment anything in images and videos. arXiv preprint arXiv:2408.00714 (2024).

[16] Isensee, F., Jaeger, P. F., Kohl, S. A., Petersen, J. & Maier-Hein, K. H. nnu-net: a self-configuring method for deep learning-based biomedical image segmentation. Nature Methods 18, 203–211 (2021).

[17] Conrad, R. & Narayan, K. Cem500k, a large-scale heterogeneous unlabeled cellular electron microscopy image dataset for deep learning. Elife 10, e65894 (2021).

[18] Xing, C., Xie, R. & Bader, G. D. Retina: Reconstruction-based pre-trained enhanced transunet for electron microscopy segmentation on the cem500k dataset. PLOS Computational Biology 21, e1013115 (2025).

[19] Heinrich, L. et al. Whole-cell organelle segmentation in volume electron microscopy. Nature 599, 141–146 (2021).

[20] Berger, D. R., Seung, H. S. & Lichtman, J. W. Vast (volume annotation and segmentation tool): efficient manual and semi-automatic labeling of large 3d image stacks. Frontiers in Neural Circuits 12, 88 (2018).

[21] Van Liedekerke, P. et al. Quantitative modeling identifies critical cell mechanics driving bile duct lumen formation. PLOS Computational Biology 18, e1009653 (2022).

[22] Van Griethuysen, J. J. et al. Computational radiomics system to decode the radiographic phenotype. Cancer Research 77, e104–e107 (2017).

[23] Quintana-Cabrera, R. & Scorrano, L. Determinants and outcomes of mitochondrial dynamics. Molecular Cell 83, 857–876 (2023).

[24] Gatti, P., Schiavon, C., Cicero, J., Manor, U. & Germain, M. Mitochondria-and ER-associated actin are required for mitochondrial fusion. Nature Communications 16, 451 (2025).

[25] Parlakgül, G. et al. Spatial mapping of hepatic er and mitochondria architecture reveals zonated remodeling in fasting and obesity. Nature Communications 15, 3982 (2024).

[26] Wang, P. T. et al. Distinct mechanisms controlling rough and smooth endoplasmic reticulum contacts with mitochondria. Journal of Cell Science 128, 2759–2765 (2015).

[27] Bonkovsky, H. L. et al. Clinical presentations and outcomes of bile duct loss caused by drugs and herbal and dietary supplements. Hepatology 65, 1267–1277 (2017).

[28] Jia, Y. et al. Atypical primary biliary cholangitis results in vanishing bile duct syndrome with cutaneous xanthomas: a case report. Diagnostic Pathology 17, 57 (2022).

[29] McConnell, M. J., Kostallari, E., Ibrahim, S. H. & Iwakiri, Y. The evolving role of liver sinusoidal endothelial cells in liver health and disease. Hepatology 78, 649–669 (2023).

[30] Ortega-Ribera, M. et al. Increased sinusoidal pressure impairs liver endothelial mechanosensing, uncovering novel biomarkers of portal hypertension. JHEP Reports 5, 100722 (2023).

[31] Amirola-Martinez, M. et al. Aspects of zone-like identity and holotomographic tracking of human stem cell-derived liver sinusoidal endothelial cells. Frontiers in Cell and Developmental Biology 13, 1528991 (2025).

[32] Marrone, G., Shah, V. H. & Gracia-Sancho, J. Sinusoidal communication in liver fibrosis and regeneration. Journal of Hepatology 65, 608–617 (2016).

[33] Gao, J., Zuo, B. & He, Y. Liver sinusoidal endothelial cells as potential drivers of liver fibrosis. Molecular Medicine Reports 29, 40 (2024).

[34] MacParland, S. A. et al. Single cell RNA sequencing of human liver reveals distinct intrahepatic macrophage populations. Nature Communications 9, 4383 (2018).

[35] Szafranska, K., Kruse, L. D., Holte, C. F., McCourt, P. & Zapotoczny, B. The whole story about fenestrations in LSEC. Frontiers in Physiology 12, 735573 (2021).

[36] Blender Online Community. Blender - a 3D modelling and rendering package. Blender Foundation, Blender Institute, Amsterdam (1994).

[37] Yushkevich, P. A., Gao, Y. & Gerig, G. ITK-SNAP: An interactive tool for semi-automatic segmentation of multi-modality biomedical images. Annual International Conference of the IEEE Engineering in Medicine and Biology Society 2016, 3342–3345 (2016).

[38] Iudin, A. et al. EMPIAR: the electron microscopy public image archive. Nucleic Acids Research 51, D1503–D1511 (2023).

