## Supplementary figures for "Multiscale volume electron microscopy of the human liver maps vascular-cellular architecture, organelle dynamics and inter-organelle communication"

### 1 Supplementary figures

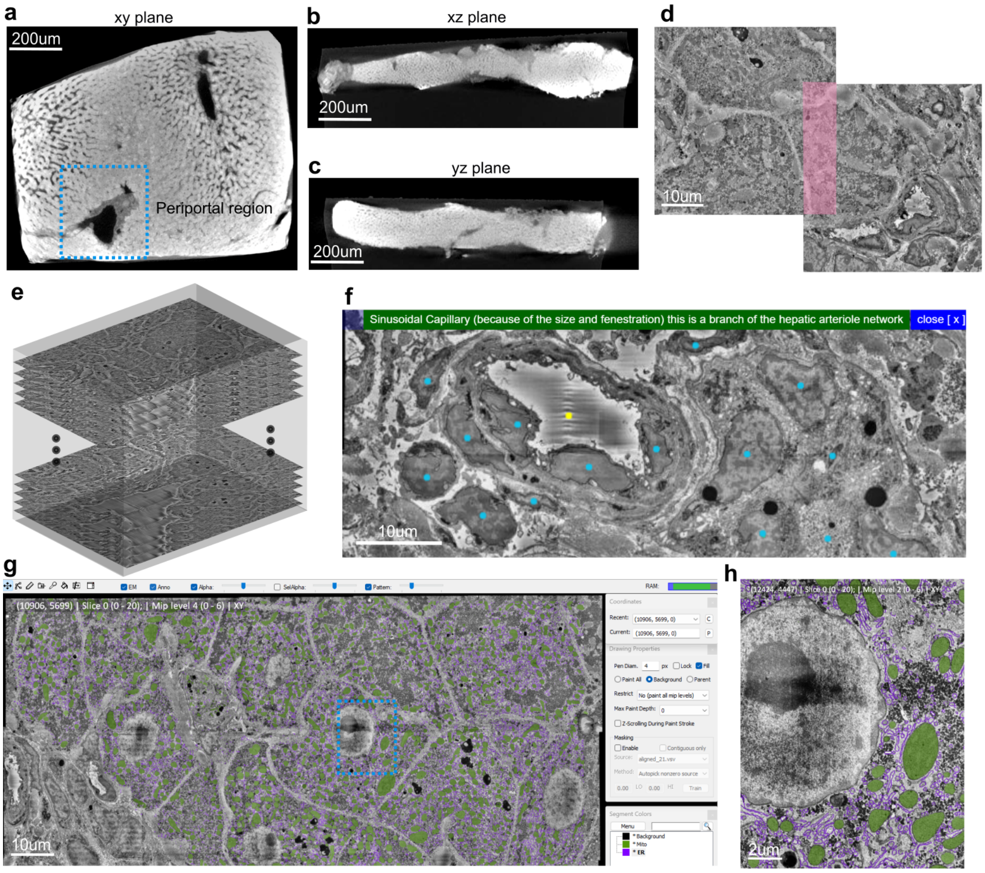


**Fig. S1 Human liver sample region of interest selection, vEM reconstruction, and manual annotation.** **a**, X-ray tomography view of the human liver tissue sample from the $xy$ plane, with the periportal region of interest highlighted (blue dashed box). **b**, X-ray tomography view from the $xz$ plane. **c**, X-ray tomography view from the $yz$ plane. **d**, Individual tiles stitched together using overlapping regions (highlighted in pink) as alignment references. **e**, Stitched sections aligned and stacked to generate a three-dimensional volume. **f**, Pathologist annotation via coordinate placement (dots) using CATMAID. Description can be shown when clicking (yellow dot). **g**, Manual segmentation performed using VAST. Mitochondria (green) and endoplasmic reticulum (purple) are shown. Blue dashed box indicates region magnified in **h**. **h**, Magnified view of the region indicated in **g**.

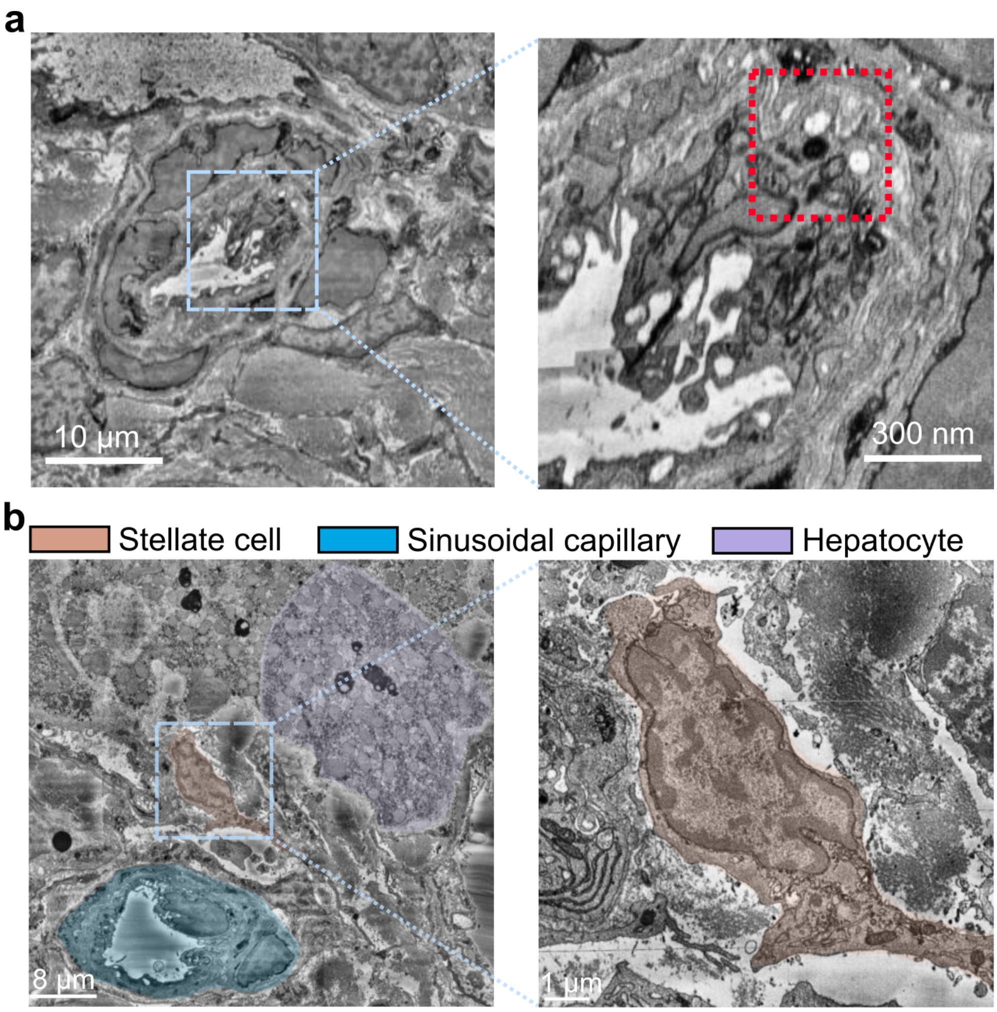


**Fig. S2 Sinusoidal capillary and stellate cell ultrastructure in the human liver volume.** **a**, Left: representative electron microscopy cross-section of the sinusoidal capillary; boxed region contains endothelial fenestrations, a defining feature of liver sinusoidal capillaries. Right: magnified view showing fenestration structures (red box). **b**, Left: representative electron microscopy cross-section showing a stellate cell located between a hepatocyte (purple) and sinusoidal capillary (blue); boxed region indicates a stellate cell (brown). Right: magnified view of the stellate cell.

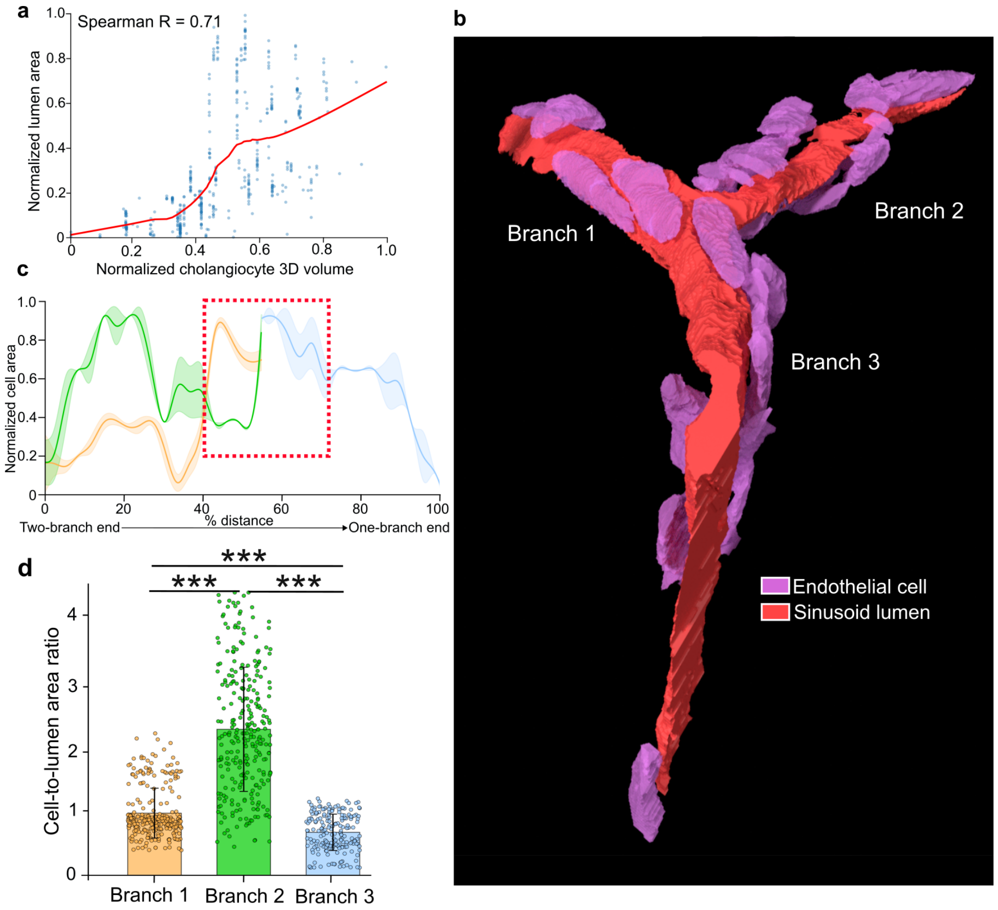


**Fig. S3 Bile duct and sinusoidal capillary structural organization in the human liver volume. a**, Correlation between bile duct lumen area and mean cholangiocyte area in three-dimensional instance-wise analysis. Red line represents locally weighted scatterplot smoothing (LOWESS) regression. **b**, Three-dimensional rendering of sinusoidal capillary lumen and surrounding endothelial cells. **c**, Endothelial cell area as a function of distance along the sinusoidal capillary, measured from the two-branch end to the single-branch end. Red dashed box indicates the region near the branching point where three branches converge. **d**, Comparison of endothelial cell-to-lumen contact ratios among sinusoidal branches (branch 1, $n = 281$; branch 2, $n = 281$; branch 3, $n = 200$ cross-sections). Statistical significance was determined by two-tailed Mann-Whitney U test with Bonferroni correction; $^{***}P < 0.001$.

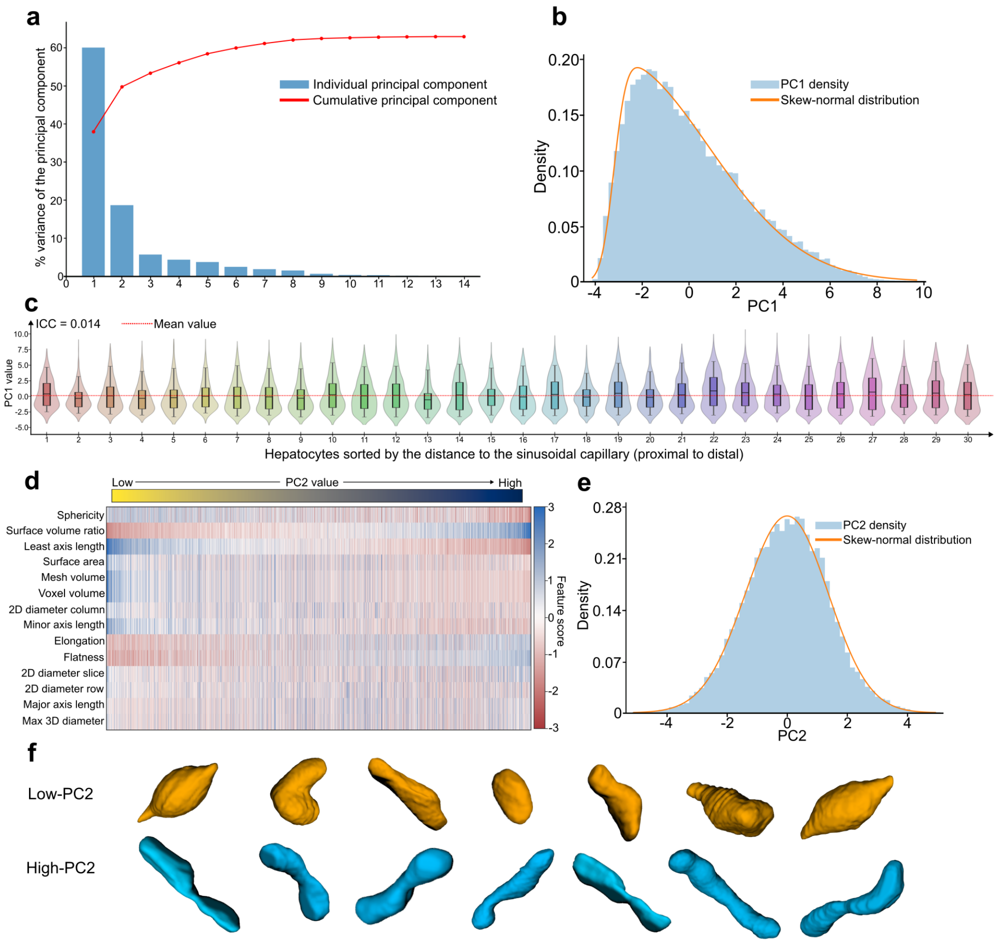


**Fig. S4 Principal component analysis of mitochondrial morphology in the human liver volume.** **a**, Scree plot showing variance explained by each principal component. **b**, Distribution of principal component 1 (PC1) values across all mitochondria, fitted with a skew-normal distribution. **c**, Violin plots showing PC1 distributions of mitochondria within 30 hepatocytes with relatively intact cell bodies. Hepatocytes are sorted by distance to the sinusoidal capillary (left to right: proximal to distal). ICC, intraclass correlation coefficient. **d**, Heatmap of morphological features across mitochondria sorted by principal component 2 (PC2) value. Each column represents an individual mitochondrion; each row represents a morphological feature. **e**, Distribution of PC2 values across all mitochondria, fitted with a skew-normal distribution. **f**, Representative three-dimensional renderings of low-PC2 and high-PC2 mitochondrial instances.

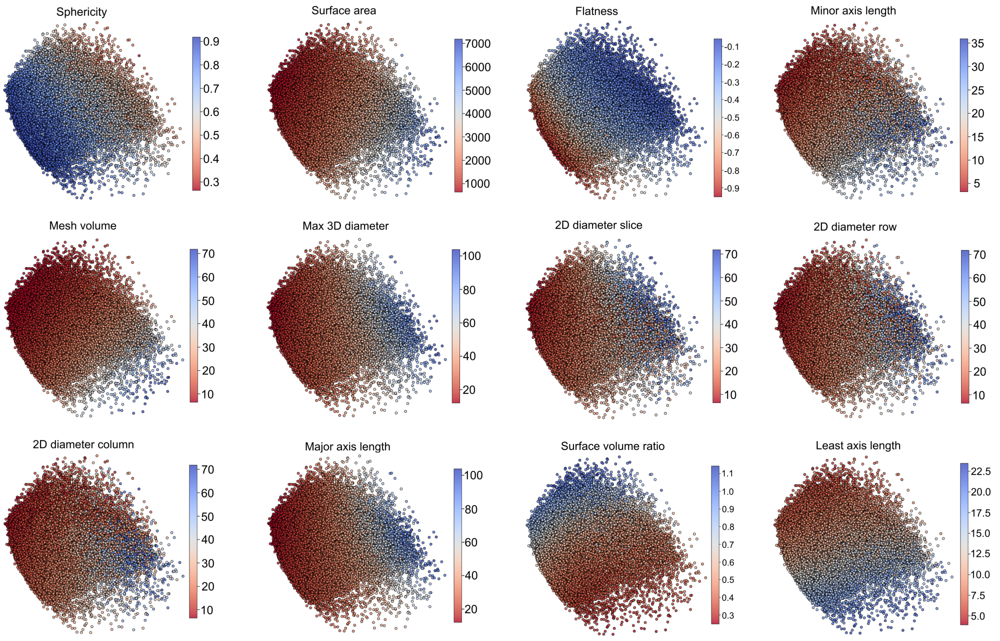


**Fig. S5 Distribution of morphological features across the PCA space.** Individual morphological features of 35,790 mitochondria projected onto the principal component analysis plot.

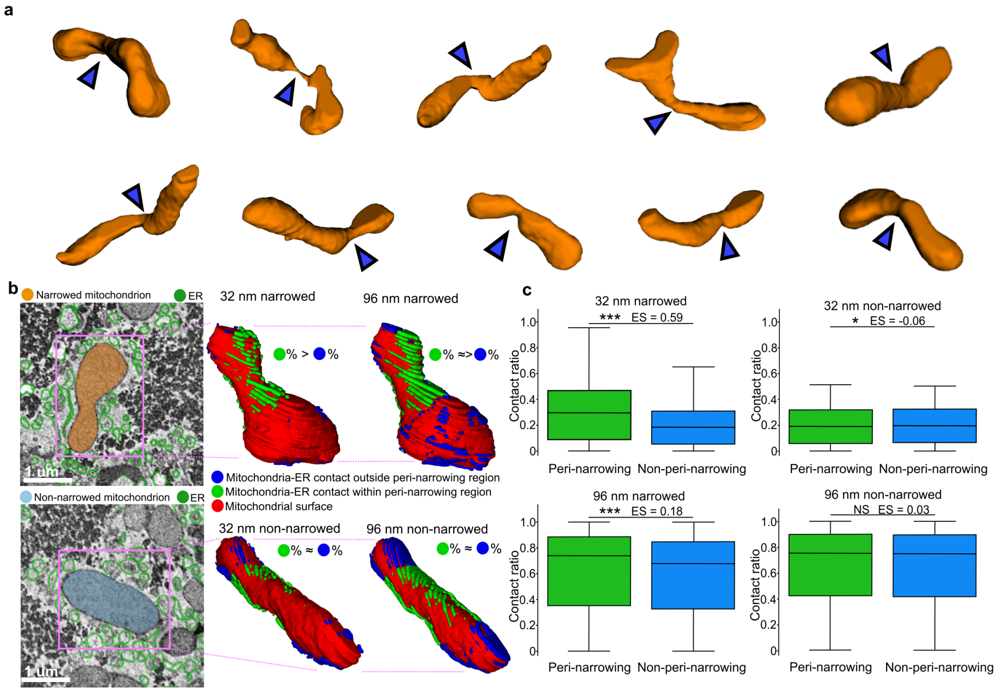


**Fig. S6 Mitochondria-endoplasmic reticulum (ER) contact preference at mitochondrial narrowing sites. a**, Three-dimensional renderings of representative high-PC2 mitochondria displaying prominent narrowing sites (blue arrows). **b**, Top row: representative narrowed mitochondrion shown in EM cross-section (left) and three-dimensional renderings illustrating ER contacts at 32 nm (middle) and 96 nm (right). Bottom row: representative non-narrowed mitochondrion shown in EM cross-section (left) and three-dimensional renderings illustrating ER contacts at 32 nm (middle) and 96 nm (right). **c**, Box plots comparing ER contact ratio between narrowed and non-narrowed mitochondria at 32 nm and 96 nm distances. Contact ratio is defined as the ER-contacted surface area divided by total mitochondrial surface area within each region. Center line, median; box bounds, 25th–75th percentiles; whiskers, minimum and maximum values. Effect size calculated by pooled median absolute deviation (narrowed, $n = 1,760$; non-narrowed, $n = 1,827$). Statistical significance determined by two-tailed Mann-Whitney U test; $*P < 0.05$, $***P < 0.001$; NS, not significant; ES, effect size.
